# Speed breeding transgenic American chestnut trees toward restoration

**DOI:** 10.1101/2025.05.19.654928

**Authors:** Thomas Klak, Hannah Pilkey, Virginia G. May, Dakota Matthews, Allison D. Oakes, Ek Han Tan, Andrew E. Newhouse

## Abstract

The American chestnut (*Castanea dentata*) was a dominant, foundational forest canopy tree in eastern North America until an imported chestnut blight (caused by *Cryphonectria parasitica*) rendered it functionally extinct across its native range. Biotechnological approaches have the potential to help restore the species, but field-based breeding advances are hampered by long generation times, ≤50% transgene inheritance, and regulatory restrictions on outdoor breeding of transgenic trees. Self-incompatibility and flowering phenology further limit generational advances and field testing of chestnuts. Our work here demonstrates that long generational times and field constraints can be circumvented by producing both male and receptive female flowers in controlled indoor environments. Additionally, we developed an embryo rescue protocol for both indoor and field conditions, in which developing embryos can be extracted and micropropagated from immature seeds between 6- and 8-weeks post pollination. These advances have enabled production of the first homozygous transgenic American chestnuts, which have produced pollen that was used for outdoor controlled pollinations and yielded nearly 100% transgene inheritance by offspring. This work also provides event-specific DNA markers to differentiate transgenic chestnut lines and identify homozygous individuals. We demonstrate that an obligate outcrossing forest tree can reach sexual maturity rapidly in controlled, indoor environments. When coupled with genomic analyses and other biotechnological advances, this procedure could facilitate the reintroduction of this iconic species.

## Introduction

The American chestnut (*Castanea dentata* [Marsh.] Borkh.) was once an ecologically, economically, and culturally significant species of eastern North American forests (United States Forest Service 1971). In the late 1800s, the invasive chestnut blight fungus (*Cryphonectria parasitica* [Murr.] Barr.) was introduced in New York and within a few decades, billions of chestnut trees succumbed to pathogen infection (Hepting 1974). Although American chestnuts are considered functionally extinct, a portion of the chestnut population survives due to the tree’s ability to asexually resprout from the root collar after blight kills it above ground (Stillwell et al. 2003). Numerous programs have been working to conserve and restore the American chestnut for over a century. One method of enhancing blight tolerance in chestnuts has been achieved through genetically engineering American chestnut via *Agrobacterium*-mediated transformation (Maynard et al. 2015). The most promising transgenic chestnut trees thus far express an oxalate oxidase (*OxO*) gene derived from wheat, which neutralizes the toxic oxalic acid delivered by the fungus (Liang et al. 2001; Zhang et al. 2013). Trees with the *OxO* transgene are collectively referred to as Darling (Powell et al. 2019), or ‘Darling #’ when referring to a specific event. Previously described Darling chestnuts include ‘Darling 4,’ a two-copy transgenic line expressing *OxO* under the *VspB* vascular promoter (Newhouse et al. 2014), and ‘Darling 54’ and ‘Darling 58,’ which are single-copy transgenic lines (Newhouse et al. 2025, Steiner et al. 2017) with OxO expressed under the CaMV35S constitutive promoter (Guilley et al. 1982).

While transgenic American chestnuts can be produced through micropropagation and acclimatization (Oakes et al. 2020), clonal propagation is not ideal for restoration goals: breeding blight-tolerant Darling trees with surviving wild chestnut trees is necessary to conserve wild-type germplasm, introgress the *OxO* gene, and increase genetic diversity (Westbrook et al. 2020). Transgenic American chestnut pollen was first used for pollinations in 2011, and ‘Darling 54’ pollen has been in use since 2016. However, progress has been hindered by the reproductive traits of the chestnut. Like many long-lived tree species, chestnuts are slow to reproduce. American chestnuts typically produce staminate (male) flowers bearing pollen in three to five years (Andrade et al. 2009), and it can take seven to ten years for trees to produce pistillate (female) flowers (Cook and Forest 1979). This can stretch to twenty or more years for trees growing in understory conditions (Paillet and Rutter 1989). Chestnuts are also largely self-incompatible, requiring outcrossing with other individual trees for successful fertilization (McKay 1942; Burnham 1988). Thus, when starting with two similar-aged field-grown seedlings, it can take upwards of seven years to intercross and produce viable offspring (Westbrook et al. 2020). Additional delays often occur when flower or seed production is reduced by external factors such as late frost, drought, insect attack, animal browse, storms, or blight infection.

When a wild-type diploid plant such as American chestnut is crossed with a compatible hemizygous (single transgene copy on one chromosome) transgenic relative, approximately 50% of progeny are expected to inherit and express the transgene. In practice, outdoor controlled pollinations using hemizygous transgenic pollen typically yield 35-45% transgene inheritance (Newhouse et al. 2025), potentially due to presence of airborne wild-type pollen during controlled pollination efforts. To generate pollen for controlled pollinations and accelerate chestnut reproduction, Darling pollen (and to a lesser extent mature Darling nuts) can be rapidly produced indoors through speed breeding.

As a plant breeding tool, speed breeding is simply the manipulation of controlled, indoor environmental and horticultural methods that accelerate the sexual maturity of plants. This process has been applied to various row crops (Ghosh et al. 2018; Watson et al. 2018), though its relevance for trees with long generation times is especially relevant. Various forms of speed breeding have been successfully applied to diverse tree including apples (Flachowsky et al. 2011; van Nocker and Gardiner 2014), *Eucalyptus* (Castro et al. 2021), citrus (Moore et al. 2016), and Douglas-fir (Kolpak et al. 2015). However, some of these rely on manipulations such as grafting or application of plant growth regulators, which may not be feasible for immature trees grown indoors.

Pioneering speed breeding work for American chestnut relied on production of male catkins, which consistently form earlier than female flowers in both artificial (indoor) and natural (outdoor) conditions. Male flowers can be induced in under one year when trees are fertilized regularly and exposed to high intensity light with a prolonged photoperiod (Baier et al. 2012; Klak et al. 2021). Typically, pollen produced indoors year-round has been collected, frozen, and used for outcrossing to wild-type trees during the outdoor breeding season. Female flowers have also been induced in under one year (Baier et al. 2012), but prior to the work presented here, crosses to female flowers rarely reached maturity. In this work, we show that mature seed production indoors is possible, in conjunction with plant care improvements such as repeated repotting as seedlings grow, fertilization of trees under high light, and control of pests and pathogens.

After outcrossing results in more distantly related individuals that have both inherited *OxO*, resulting transgenic trees can be intercrossed. After outcrossing more distantly related individuals that have both inherited OxO, the resulting transgenic offspring can themselves be intercrossed. Crossing two hemizygous trees will theoretically result in a pool of offspring that is approximately 25% non-transgenic, 50% single-copy (inheriting *OxO* from one parent), and 25% two-copy (inheriting *OxO* from both parents). Darling trees with two copies of *OxO* are considered homozygous for the transgene. Pollen from trees homozygous for *OxO* would be ideal for producing Darling seeds from controlled pollinations, since 100% of offspring from crosses with Darling homozygous pollen are expected to inherit the *OxO* gene for blight tolerance. Trees homozygous for *OxO* are not a restoration priority, as they do not necessarily provide environmental benefits beyond those present in hemizygous trees. However, *OxO* homozygosity would be valuable for research and consistent production of increasingly diverse blight-tolerant offspring, especially from outcrosses to mother trees with few flowers or those that are declining from blight. Molecular methods to distinguish homozygous from hemizygous transgenic Darling chestnut trees have not been previously published.

Chestnuts can also be initiated from various tissues and micropropagated using tissue culture to contribute to the breeding program. Through tissue culture, we have produced flowers on clonally propagated plantlets, and flowers which mature into viable nuts on seedlings. To protect limited and valuable chestnuts, collection of immature embryos (here termed “embryo rescue” following Pilkey [2021]) and *in vitro* germination could help indoor-pollinated chestnuts reach maturity. Embryos are a common explant for initiating various plant species in tissue culture, including walnut (*Juglans* sp.) (Kaur et al. 2006), Japanese holly (*Ilex crenata*) (Yang et al. 2015), and grape (*Vitis* sp.) (Jiao et al. 2018). Because of their juvenility, embryos from many plant species are inclined to regenerate and multiply *in vitro* with relative ease (Kyte et al. 2013). To our knowledge, embryo rescue has not been previously reported with American chestnut.

The objectives of this study were to 1) describe production of indoor American chestnut pollen and female flowers that are capable of producing fertile nuts, 2) test initiation timing of an *in vitro* embryo rescue protocol for American chestnut, 3) test the *in vitro* embryo rescue protocol with embryos produced through indoor pollinations and, 4) develop two distinct molecular methods for distinguishing homozygous from hemizygous trees, including a copy number test using qPCR and a multiplex PCR genotyping assay.

## Methods

### Transgenic American chestnut speed breeding conditions

Speed breeding methods were deployed to produce quantities of pollen and sexually mature chestnuts under controlled indoor conditions by eliminating seasonality for this deciduous species. While we have collectively observed early flowering hundreds of times under diverse indoor high-light conditions, our experience identified seven crucial factors for effective chestnut speed breeding. Comparable material with similar specifications may be used based on the materials listed herein. (1) Full-spectrum grow lights (Mars Hydro SP 3000) on 15-16 hour photoperiods year-round, with PAR levels at the top seedling leaves near the outdoor June 21 noon maximum (approximately 600 to 1000 µM/m^2^/s at upper leaves). (2) Acidic, porous potting mix (Premier Pro-Mix BK55), enhanced with micronutrients (Micromax G90505 1.3 mL/L), time-release granular fertilizer (Osmocote Plus 1.3 mL/L), and perlite for additional aeration. (3) Water-soluble fertilizers administered every two to three days, ideally including a flowering nutrient (Roots Organic Terp Tea Bloom 4 mL/L) and micronutrients (alternating FloraFlex, Drip or Mills A&B [8 mL/L], with amendments of phosphorus, potassium [Drip flex 2.6 mL/L] and calcium [Drip CalMag 1.3 mL/L]). (4) Larger pot sizes as each plant grows, to confine but not fully restrict roots. Nuts with exposed radicles were sown in one- or two-gallon pots, and successively transplanted to 5–7 gallon pots and then to as large as 15 gallon pots before they became root-bound. (5) Growth room temperatures ideally in the range of 24°C daytime and 17°C at night, averaging approximately 20°C. (6) Relative humidity controlled to approximately 30-50%. (7) Timely identification and treatment of pest and pathogen outbreaks as part of an integrated pest management scheme, including treatments for fungus gnats, spider mites, and powdery mildew.

### Indoor production of pollen and fertile nuts on transgenic American chestnut seedlings

We typically collect transgenic pollen from catkins directly onto glass microscope slides, which can be used immediately for pollinations or stored for freezing in air-tight slide fixing jars (e.g., Slide-Fix Caplugs, Evergreen Labware). Receptive female flowers were typically pollinated multiple times with the same pollen to maximize chances of pollination during optimal receptivity. Compared to outdoor breeding, the indoor setting with smaller trees allowed easier access to both female flowers and pollen and facilitated multiple pollinations. Additionally, indoor pollination was performed throughout the year, in contrast to the approximately two-week pollination window outdoors. After successful pollination, fruit development (bur growth) was observed and visibly mature burs were confirmed to contain fertile seeds.

We observed that lapses, such as high temperatures or low relative humidity, could cause the sensitive female flowers to abort. Despite this sensitivity, we observed more than half of all lab-generated female flowers reached maturity over a recent two-year period. Burs began to split and fully mature chestnuts were harvested indoors on average 13 weeks after the first pollination. Chestnuts were then stored for approximately two months at room temperature in sphagnum moss moistened with Bt (e.g. Summit Mosquito Bits, to suppress fungus gnats). For comparison, typical stratification recommendations for chestnuts specify approximately 2-3 months at 1-4.5°C (Growing Chestnuts 2025). Nuts with radicles visible after this shortened stratification were then sown in two-gallon pots and gradually introduced to increasing amounts of fertilizer and intensity and day length under grow lights.

### Embryo rescue from immature American chestnut seeds

Wild-type seeds were used to test the *in vitro* embryo rescue method. Two wild-type flowering American chestnut trees in Sherburne, NY were selected to be pollinated. To prevent open pollination and ensure accurate timing, immature pistillate flowers were bagged with standard 3D pollination bags (PBS International, York, United Kingdom). When female flowers became receptive, wild-type pollen was applied to flowers in 18 pollination bags on each tree. One pollination bag on each tree was left unopened to serve as a non-pollinated control. Pollination bags covered the flowers for the duration of the study to prevent open pollination and provide protection. The pollination bags containing flowers were cut from the trees at intervals of 2 weeks, 4 weeks, 6 weeks, and 8 weeks after the controlled pollinations were conducted. Two pollination bags were removed from each tree for a total of four pollination bags for extractions every two weeks. At eight weeks the non-pollinated control bags were cut from both trees.

To sterilize the tissue prior to micropropagation, the immature chestnuts were first excised from the involucre (developing bur). Chestnuts were sprayed with 70% ethanol for 30 seconds, then submerged and continuously shaken in a solution of 50% bleach (~3% sodium hypochlorite) plus 0.01% Tween-20 for 15 minutes, followed by three, five-minute rinses with sterile distilled water. To remove the seed coat, the seed was cut latitudinally at the hilum and a longitudinal cut was made to open and peel away the pericarp. Once the seed coat was removed, ovules or embryos were aseptically removed. Explants were considered an ovule if the fertilized ovule could not be distinguished from its unfertilized neighbors. The explant type was considered an embryo if the functional ovule (FO) that had been fertilized could be identified. All observable ovules or embryos were extracted and placed with the micropyle facing downward into a multiplication medium (full-strength Lloyd and McCown Woody Plant medium salts, 109 mg/L Nitsch and Nitsch vitamins, 1 µM benzyladenine, 0.01 µM indole-3-butyric acid, 3% sucrose, 0.7% agar, with pH adjusted to 5.5). Explants were kept on a light shelf receiving 31 µmol/m^2^/s of light with a 16-hour photoperiod. All ovules and embryos were transferred to fresh media every two weeks.

Explants were observed for germination every two weeks at the end of their transfer cycle. In this experiment, germination was defined as the emergence of a shoot apical meristem. All explants were given a value of “0”. When an embryo germinated, it was given a value of “1”. Data were analyzed using R version 3.4.3. ANOVA analyses were used for germination rates, and Tukey’s Honest Significant Difference was used to determine the significance between treatments. Significance was evaluated at α=0.05. Explants were also monitored for contamination and death. At the end of the experiment, the overall germination rate and elapsed time until germination were observed. Contamination rates for each treatment were calculated, however, explants that were contaminated were removed from the germination rate calculation. Shoot maintenance (including multiplication and elongation stages) in tissue culture, as well as *ex vitro* rooting and potting protocols were performed using previously established methods for American chestnut (Oakes et al., 2020).

### DNA isolation

50-100 mg of fresh, young chestnut leaf tissue from plants grown indoors were manually disrupted in a 1.5 mL Eppendorf tube and extracted using DNeasy Plant Pro kit (Qiagen 69204) using the standard manufacturer protocol.

### Reconstruction of ‘Darling 54’ T-DNA insertion junctions

Illumina reads from 16020, one of the T1 founder lines used for Darling chestnut breeding were generated by the HudsonAlpha Genome Sequencing Center and mapped to T-DNA using Geneious (11.1.5) program using the short read paired-end setting. Next, paired reads spanning right border (RB) and left border (LB) were manually extracted and assembled using SPAdes assembler (3.10.0) (Bankevich et al. 2012). Contigs from RB and LB reads were both mapped to both T-DNA and Ellis v1.1 genome assembly (Westbrook et al. 2025) using BLAST to determine T-DNA junction to the Ellis v1.1 Chr04. Reconstruction of the intact, single copy T-DNA insertion in ‘Darling 54’ is presented in Supplemental Figure 1.

### Real time quantitative PCR assays

Quantitative PCR reactions used Bio-Rad iTaq Universal SYBR Green Supermix in 20µl reactions including one ng template DNA and one µM final concentration of each primer. Each biological sample was done in triplicate for each primer set (*OxO* as described below, with *actin* and *ubiquitin* reference genes; see Supplemental Table 1 and Carlson et al. (2022)). Amplification was carried out in a Bio-Rad CFX Connect qPCR cycler. The qPCR program began with a three-minute denaturation step at 95°C, followed by 39 cycles of 95°C for 10s, and 60°C for 30s with reads taken at the end of each 60°C step. This was followed by a melt curve (65°C to 95°C in 0.5°C increments) to check for single specific PCR products and specificity of amplification. Copy number controls were ‘Darling 58’ (known single copy) and ‘Darling 4’ (known two copy). Relative copy number values were analyzed via normalized expression (ΔΔCq) using Bio-Rad CFX Maestro software.

### Genotyping assays

*OxO* qPCR primers were described previously (Zhang et al. 2013) but were used in our experiments for end-point PCR detection of the *OxO* T-DNA construct. The ‘Darling 58’ event was described in the USDA-APHIS Petition for nonregulated status and supplemental information (Newhouse et al. 2025). For ‘Darling 54,’ RB and LB PCR primers were designed based on reconstructed T-DNA junctions and multiplex PCR was designed based on the deletion on Caden.04G062600. Primer sequences and expected PCR sizes are provided in Supplemental Table 1. PCR was performed using 2x GoTaq Green Mastermix (Promega M7122). Approximately two ng of DNA were used per 20 µL PCR reactions containing two µM of each primer. All reactions were run with 40 cycles of 94°C for 30s, 55°C annealing for 30s and 68°C extension for 60s. Final PCR products were run on a 1.5% TAE gel and imaged after ethidium bromide staining.

### Homozygous transgenic plant development and pollen production

Since American chestnuts are generally self-incompatible, and all transgenic chestnuts of a given event are derived from a single founder genotype, genetic diversification requires outcrossing transgenic individuals over multiple generations. Intercrosses between transgenic lines were performed when both pollen and female flowers were available on T2, T3 or T4 transgenic trees that were not full siblings. Some immature seeds were subjected to embryo rescue protocols to speed recovery of offspring and are represented by the TGA, TGG and TGJ families in this report. For the ULL family from which we obtained fully mature burs indoors, attention to each of the speed breeding parameters was crucial. In the case of the TGA family, ‘Darling 54’ T2 pollen was crossed to T2 individuals derived from different families to minimize relatedness. After female flowers were observed on indoor potted transgenic plants, transgenic pollen was applied as described above. As the fertilized burs developed, they were harvested via embryo rescue. Embryos were germinated *in vitro*, propagated, and advanced to shoot culture. Transgene homozygosity of the rescued embryo cultures was confirmed by both qPCR for copy number and T-DNA genotyping using leaf tissue. One of the confirmed homozygous embryo cultures (TGA002) was propagated, rooted, acclimatized, and maintained under the same high-light conditions described above until catkins developed. Pollen from TGA002 was first produced under high light in the winter of 2023/2024 and stored at −80ºC to preserve pollen viability.

### Analysis of Mendelian ratio of D54 crosses

The analysis of Mendelian inheritance ratios from embryo rescue and mature seeds were treated as independent populations in this analysis using the families derived from crosses between two hemizygous D54 parents. Using genotype calls obtained from D54 multiplex PCR, a table was constructed with the observed (actual) genotype calls based on D54 multiplex PCR followed by the expected 1:2:1 wild-type:hemizygous:homozygous genotypes in Microsoft Excel (Version 16.79.2). The function ‘CHISQ.TEST(actual_range, expected_range)’ was then used to generate the expected p-values from the observed (actual) and expected ranges on the Excel sheet.

### Outdoor controlled pollinations with homozygous transgenic pollen and early testing

In the summer of 2024, 15 wild-type chestnut mother trees at four USDA-APHIS permitted field locations in New York state were selected for controlled pollinations using pollen homozygous for *OxO* (TGA002). In mid-June, immature pistillate flowers on the wild-type mother trees were bagged to prevent open pollination. Approximately two weeks later, homozygous pollen (TGA002) was applied to receptive stigmas and pollination bags were placed back over the developing flowers. On one of the mother trees (#1 in Table 1), pollen hemizygous for *OxO* (UET001) was also applied to additional female flowers as a control.

**Table 1.** Inheritance rates of the OxO transgene in offspring of controlled crosses with both hemizygous and homozygous pollen. Testing time (early vs. mature) is shown in weeks after pollination; Mother Tree numbers refer to separate individual trees. Mother tree 4* is a Japanese/European hybrid ‘Bouche de Betizac’; all others are fully American to the best of our knowledge.

| Pollen type | Mother Tree | Early testing<br>(8 weeks) |  | Mature seed testing<br>(~14 weeks) |  | Mean OxO<br>Inheritance<br>Rate |
| --- | --- | --- | --- | --- | --- | --- |
|  |  | n Positives /<br>n Tested | OxO<br>Inheritance<br>Rate | n Positives /<br>n Tested | OxO<br>Inheritance<br>Rate |  |
| **Hemizygous**<br>**(UET002)** | 1 | 17/47 | 0.36 | 102/224 | 0.46 | **0.41**<br>**(n=271)** |
| **Homozygous**<br>**(TGA002)** | 1 | 41/41 | 1 | 35/35 | 1 | **0.95**<br>**(n=578)** |
|  | 2 | 15/15 | 1 | 17/22 | 0.77 |  |
|  | 3 | 12/13 | 0.92 | 3/3 | 1 |  |
|  | 4* | 63/63 | 1 | 22/30 | 0.73 |  |
|  | 5 |  |  | 54/54 | 1 |  |
|  | 6 |  |  | 24/26 | 0.92 |  |
|  | 7 |  |  | 125/131 | 0.95 |  |
|  | 8 |  |  | 6/6 | 1 |  |
|  | 9 |  |  | 7/7 | 1 |  |
|  | 10 |  |  | 1/1 | 1 |  |
|  | 11 |  |  | 2/2 | 1 |  |
|  | 12 |  |  | 5/5 | 1 |  |
|  | 13 |  |  | 69/74 | 0.93 |  |
|  | 14 |  |  | 47/50 | 0.94 |  |

At eight weeks post-pollination, a portion of developing seeds were removed from four of the 15 mother trees (1 – 4 on Table 1) for early *OxO* inheritance testing. Several open-pollinated seeds from each mother tree were also collected to serve as wild-type controls. Immature seeds were peeled from the involucre. All fertilized seeds containing an embryo were peeled, and tissue from the embryo was sliced into segments and collected into microcentrifuge tubes. Embryos were tested for presence and activity of the OxO enzyme using a colorimetric histochemical assay regularly deployed on other chestnut tissues (Liang et al. 2001). All other TGA002 control-pollinated flowers from the 15 mother trees were harvested at maturity (~14-16 weeks post-pollination). Cores from the cotyledons of mature, viable seeds were tested for OxO inheritance and activity using the same histochemical assay.

## Results

### Indoor production of pollen and female flowers, bred to produce fertile nuts

An overview of our speed breeding processes, including induced female and male flowers from transgenic American chestnuts, is shown in Figure 1. Owing to the shortened maturation time frame from speed breeding, pollen-bearing male catkins were observed as early as five months after sowing. Most seedlings produced pollen within one to two years after sowing. The emergence of bisexual catkins including female flowers ranged from seven months to more than two years after sowing. Intercrosses between two hemizygous transgenic seedlings can potentially give rise to the three genotypic outcomes indicated, including homozygous transgenic lines. On the bottom left of Figure 1, female flowers that were successfully fertilized are able to develop into full-sized burs that remain green, which can be harvested to obtain fully mature seeds in about 13 weeks. In conjunction with the modified stratification described above, this process resulted in more rapid radicle emergence (as little as one month), and resulting seedlings could be transplanted directly in soil in about two months after harvest. These seedlings can then be subjected to the next speed breeding cycle, repeating the process and reducing the seed-to-seed time to just under two years.

**Figure 1.**
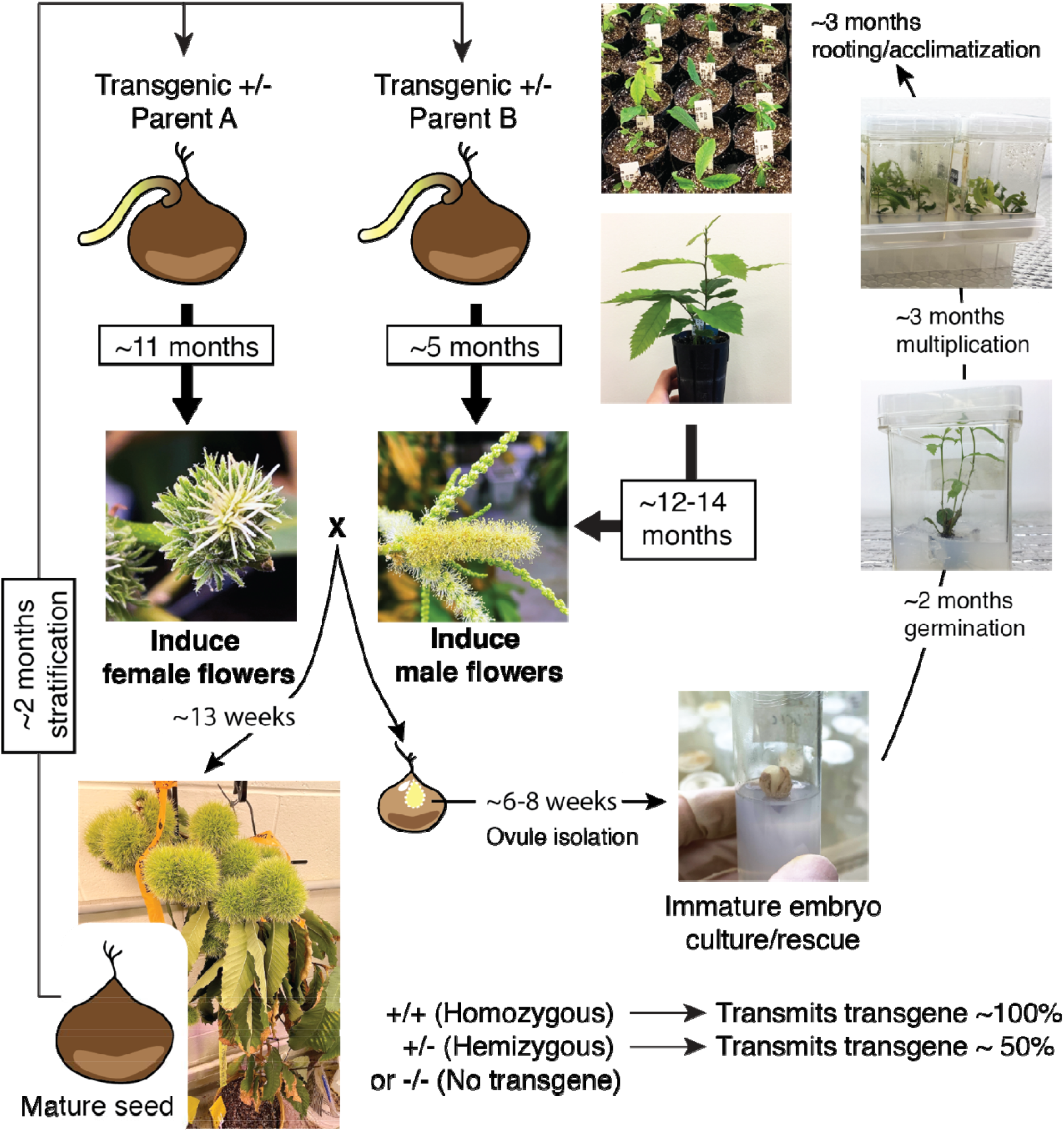
Transgenic American chestnut speed breeding scheme. Speed breeding of transgenic American chestnut can result in the induction of male flowers in as little as five months and female flowers in as little as 11 months after seed stratification. Some of the crosses performed result in fully matured burs (left downward arrow). Mature seeds can be harvested from fully matured burs and, after two months of stratification, seedlings can be cycled back for indoor flower production. Premature burs can be subject to immature embryo rescue via tissue culture (right downward arrow). Manually excised functional ovules were surface-sterilized, germinated and multiplied in vitro, and rooted plantlets on soil cycled back for indoor flower production. Genotyping of transgenic seedlings from either mature seeds or embryo rescue can be performed to assess the copy number of the transgene of interest. Bolder arrows denote seedling or plantlet subject to speed breeding flower production, while lighter arrows denote timelines for each stage. Notations: −/−indicates no transgene, +/−indicates a hemizygous transgenic line, and +/+ indicates homozygous transgenic line.

Immature burs that are not fully expanded but remain attached for at least six weeks are also viable for speed breeding with the use of embryo rescue methods as shown on the bottom right of Figure 1. In some cases, burs aborted prematurely, but ovules from immature seeds were embryo-rescued, followed by *in vitro* germination and multiplication, which takes around five months. Acclimatization and rooting of the multiplied lines take around three months, but can yield many clonal plantlets at this stage. In our experience, rooted embryo rescue lines take 12-14 months to produce male flowers (pollen) under speed breeding conditions.

### Initiation timing of *in vitro* embryo rescue for American chestnut

Embryo rescue tests on immature burs from two wild-type American chestnuts helped determine the age and conditions necessary for successful germination of dissected ovules. As ovules mature, they are called embryos; both terms are used here with a transition at approximately six weeks. A total of 100 chestnuts were removed from immature burs on involucres from controlled pollinated chestnuts at various time points (Figure 2). Between one and 15 healthy ovules were excised from each chestnut and put into growth medium. In total, 363 healthy ovules or embryos were excised from controlled pollinated chestnuts (2-weeks: 18 chestnuts [n=131 ovules]; 4-weeks: 29 chestnuts [n=117 ovules]; 6-weeks: 40 chestnuts [n=93 ovules or embryos]; 8-weeks: 18 chestnuts [n=17 embryos]). Significant differences between extraction time treatments were observed in mean germination rate (F=11.26, p<0.001) and contamination rate (F=10.66, p<0.001) (Figure 2A). The mean germination rate of ovules or embryos extracted from each treatment was: 0% at two weeks, 2% at four weeks, 8% at six weeks, and 23% at eight weeks. Contamination rates in treatment were: 0% at two weeks, 5% at four weeks, 14% at six weeks, and 45% at eight weeks. The development and differentiation of ovules into zygotic embryos can be observed by eight weeks post-pollination (Figure 2B). Average time elapsed between embryo rescue and *in vitro* germination was 133 days for four-week, 114 days for the six-week, and 70 days for the eight-week treatment. Embryos from the eight-week time point germinated significantly faster than four- and six-week time points (F=3.982, p=0.004).

**Figure 2.**
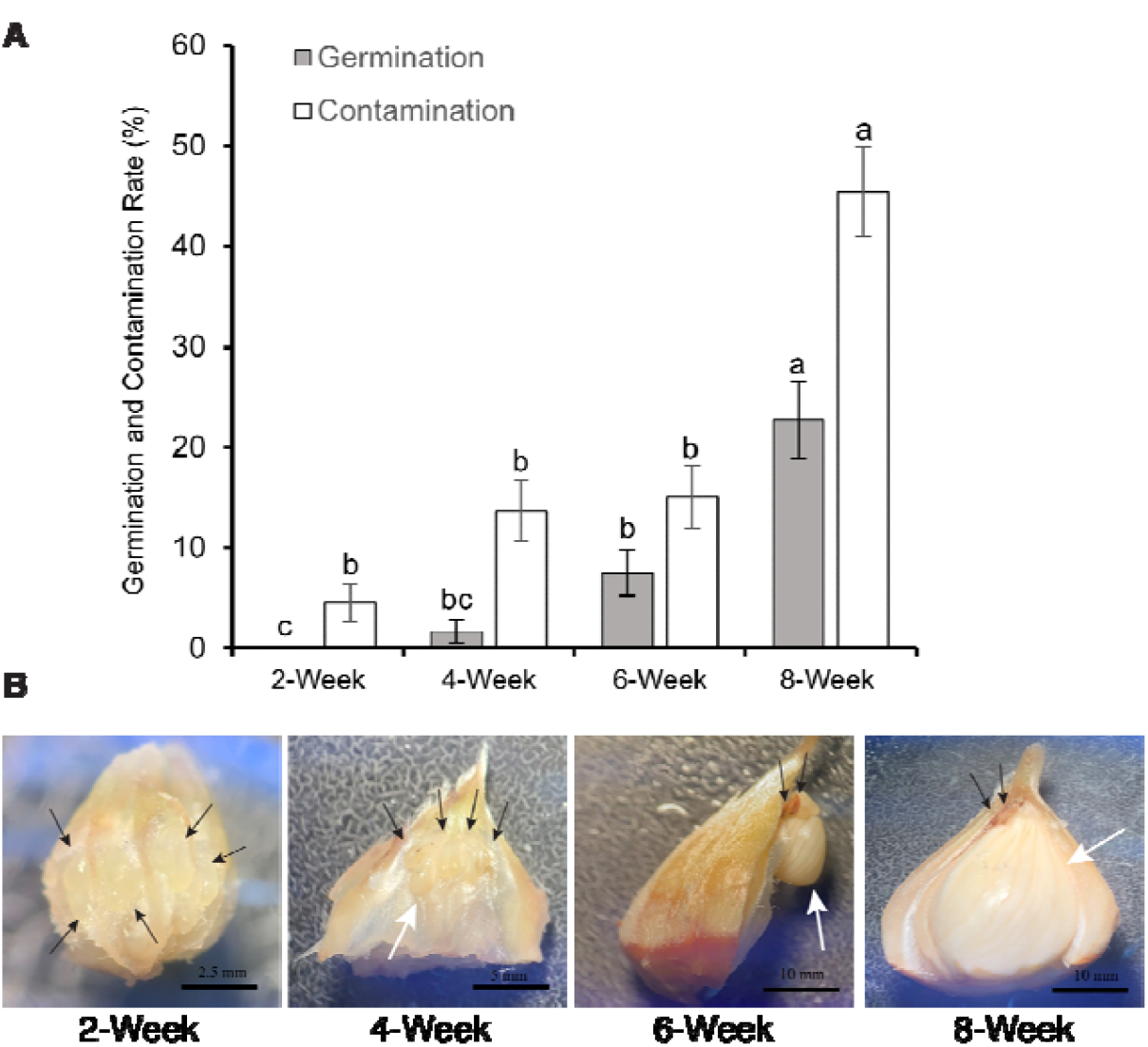
Timing of embryo rescue in American chestnut immature embryos. (A) Germination and contamination rate from American chestnut ovules collected 2-, 4-, 6- and 8-weeks after pollination. Error bars indicate +/−one standard error of the mean. (B) Photos of corresponding stages dissected from each week. In each inset image, black arrows point to immature ovules and white arrows point to functional ovules.

### Test of the *in vitro* embryo rescue protocol and mature seed viability from indoor pollinations

Indoor controlled pollinations between receptive female (ULL family) and pollen-producing Darling American chestnut lines (T2, T3, and T4) led to a total 33 burs, of which 18 were harvested while immature and 15 matured on the plant. From the 18 immature burs (six to eight weeks old), 44 ovules were dissected and subjected to embryo rescue, resulting in 24 embryo rescue individuals that germinated *in vitro* and were tested for *OxO* copy number. These individuals represented multiple offspring generations and were derived from several families (Supplemental Table 2).

From the 15 fully mature burs produced indoors, 37 fertile nuts were obtained, germinated and grown in soil. In the speed breeding setting, we obtained on average, 2.5 full-term, viable nuts per bur.

### Molecular methods for distinguishing homozygous from hemizygous transgenic American chestnut trees

We have previously demonstrated that transgene copy number (and therefore zygosity) can successfully be determined on transgenic chestnut plants using real time quantitative PCR (qPCR) (Newhouse et al. 2014; Newhouse et al. 2025). The bar plot in Figure 3A based on qPCR data indicates two copies of *OxO* in ‘Darling 4,’ a known two-copy T0 transformant (Newhouse et al. 2014), zero copies in the Ellis wild type, and one copy in SX58 (T0 of ‘Darling 58’) as expected. This method also correctly identified a homozygous Darling line TGA002 with two *OxO* gene copies, and two hemizygous Darling lines, TGG001 and TGJ001, that show a single copy of the *OxO* gene (Figure 3A). However, this qPCR assay only targets a portion of the *OxO* DNA sequence, and does not take the transgene insertion location into account. Based on the T-DNA insertion sites of ‘Darling 54’ and ‘Darling 58,’ we designed simplex and multiplex genotyping assays (Figure 3B) to test for the endogenous insert position and differentiate D54 versus D58 transgene location. D58 T-DNA border primer sets successfully amplified from SX58, while D54 T-DNA border primer sets amplified from the ULL and TGG breeding lines. The multiplex primer sets were designed to provide T-DNA homozygosity over the insertion site and work as expected, as described in detail in the caption of Figure 3B. All Darling material currently used for pollen production is derived from the D54 T0 line, SX54, though some was previously misidentified as D58 (Steiner et al. 2017; Newhouse et al. 2025).

**Figure 3.**
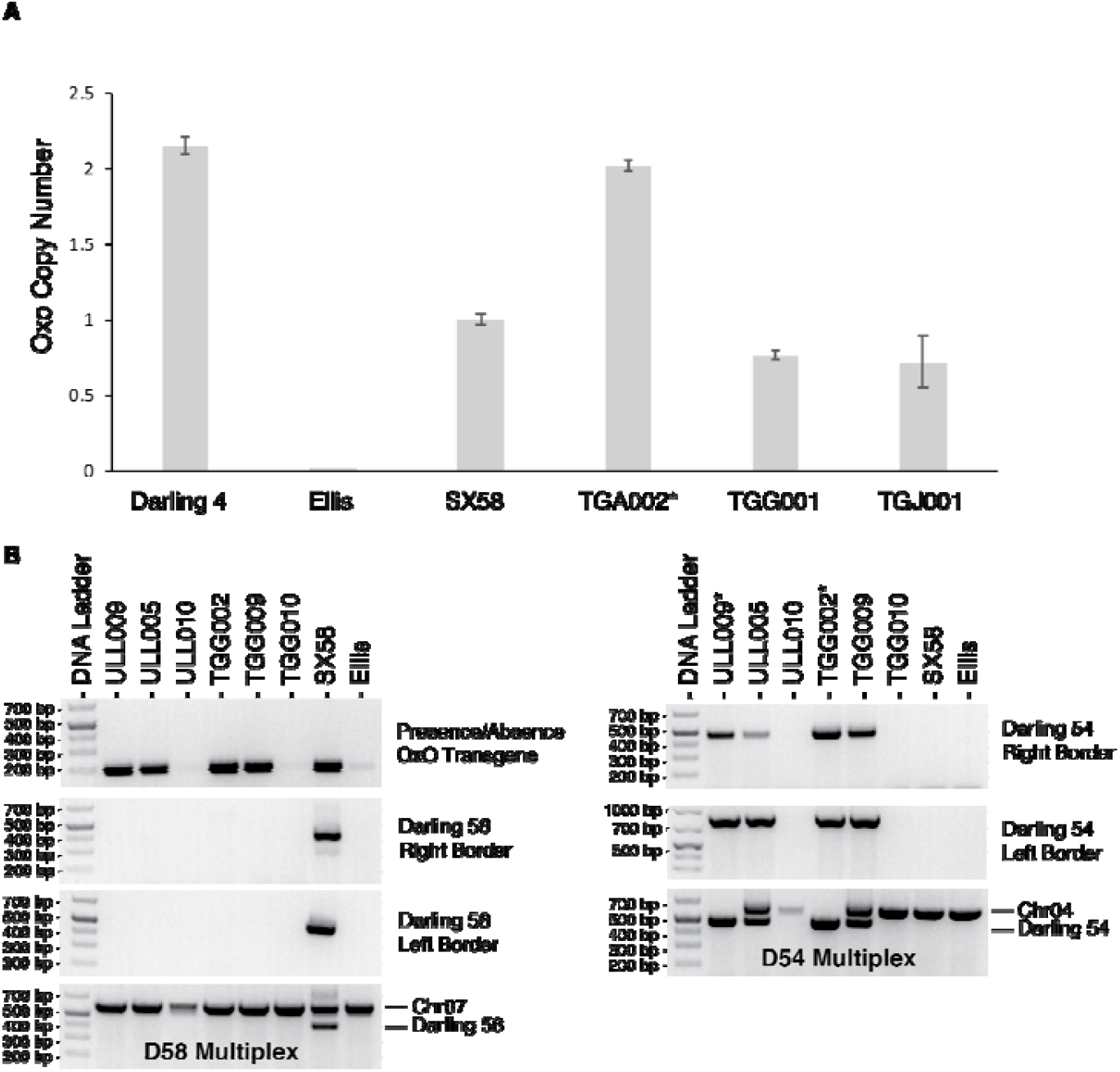
Genotyping methodologies for ‘Darling 54’ and ‘Darling 58’ from embryo rescues and seedlings from mature nuts. **(A)** Bar plots of the relative OxO copy number from quantitative PCR, showing two-copy T-DNA present in T0 ‘Darling 4’ and absence/zero-copy of OxO-containing T-DNA in the wildtype Ellis line. T0 ‘Darling 58’ (SX58) shows a single-copy state, T3 TGA002 shows two-copy state (*indicates a potentially homozygous line based on qPCR), T5 TGG001 and T5 TGJ001 show single-copy states. (**B**) Genotyping assay for OxO, ‘Darling 58’ (D58) and ‘Darling 54’ (D54) on two T5 transgenic families (ULL and TGG), T0 ‘Darling 58’ (SX58) and the Darling wild type progenitor (Ellis). The OxO qPCR primer set determines the presence or absence of the OxO transgene regardless of insert location. D58 RB and LB determine presence of the OxO T-DNA on Chr07, while D58 Multiplex provides amplification of the insertion site on Chr07 and D58 LB. Similarly, D54 RB and LB determine presence of OxO T-DNA on Chr04, while D54 Multiplex provides amplification of the insertion site on Chr04 and D54 RB. Based on these assays, ULL and TGG T5 families are derived from D54. ULL009 and TGG002 are homozygous for D54 while ULL005 and TGG009 are hemizygous for D54. ULL010 and TGG010 are wild-type null segregants.

In total, three of the 24 lines derived from embryo rescue and eight of the 99 mature fertile nuts we genotyped were homozygous for *OxO* (Supplemental Table 3). We successfully recovered all three genotypic outcomes (non-transgenic, hemizygous, and homozygous) in seedlings from embryo rescued as well as fertile nuts.

### Outdoor pollinations with homozygous pollen

Outdoor controlled pollinations with homozygous ‘Darling 54’ pollen yielded a total of 578 seeds in 2024, of which 132 were harvested for early testing and 446 were harvested at maturity. Concurrent pollinations with hemizygous pollen on one of the same mother trees yielded 47 seeds harvested early and 224 harvested at maturity. Of the early harvests from a homozygous pollen parent, all but one seed (131/132) tested positive for *OxO*, while mature harvests showed 417 *OxO*-positive out of 446 total nuts, for a combined total of 95% *OxO* inheritance from a homozygous parent (Table 1). In contrast, pollinations with hemizygous pollen resulted in a combined total of 41% inheritance (119 out of 271 total harvested nuts), which is typical based on our extensive experience with transgenic pollinations.

## Discussion

This paper reports on three novel contributions to American chestnut restoration that also have potential applicability to other *Castanea* species and hybrids as well as to other tree species. First, we have bred fully mature transgenic chestnuts indoors. Most previously described speed breeding methods are aimed at rapidly advancing generations for domesticated crop species (Ghosh et al. 2018; Watson et al. 2018; Bhatta et al. 2021). By adapting speed breeding techniques for trees, we show that American chestnuts can be induced to produce both pollen and viable female flowers, bypassing the temporal constraints of the annual outdoor growing season, and in a much-reduced maturation timeline compared to outdoor breeding. In turn, male- and female-mature chestnut plants can be strategically and successfully crossed indoors to advance chestnut generations. These contribute to overcoming the founder effect of a single transgenic event and to diversifying the genetic pool for potential restoration plantings (Newhouse and Powell 2021; Westbrook et al. 2021). Our methods to speed breed American chestnut can be replicated in almost any lab or growth room using cost-effective, space-advantageous and widely available wide-spectrum LED lights provided that the right cultural conditions are met. The parameters may also have to be refined at different locations, depending on the environmental controls available in each speed breeding lab. We also showed that fully developed mature burs containing fertile nuts can also be produced and harvested under speed breeding conditions. Our initial yield of 2.5 nuts per bur is consistent with yields from outdoor controlled crosses with transgenic pollen. Stratification can be shortened to around two months (until they grow a substantial radicle) before sowing in soil. After a few weeks of acclimatization, newly sprouted seedlings that survive can be genotyped, and cycled back into the speed breeding process.

Second, the embryo rescue methods and results reported in this paper are novel to American chestnut. The most suitable time point to extract functional ovules from American chestnut for embryo rescue is at six to eight weeks after pollination, though some germination was observed as early as four weeks. Dissected functional ovules typically take two to four weeks to germinate. Multiplication and rooting of germinated embryos can also result in rapid production of whole plants that can be used for speed breeding or other field studies. This also allows the dissection of seeds from immature burs from speed breeding that would otherwise have aborted early, providing a way to use this intentionally or as emergency rescue of important crosses. Embryo rescue feeds back into the speed breeding cycle (Figure 1), by making it possible to produce pollen from a plant that is homozygous, or a tree that performs well in the field, without the delay associated with traditional breeding to produce mature nuts. Another advantage of working with immature burs for embryo rescue is that the embryos can be cloned easily (Xing et al. 1999; McGuigan et al. 2020), which allows for larger population increases in a shorter time for genotypes of interest. However, seedlings transferred from tissue culture to soil may take longer to flower compared to seedlings from mature seed (Figure 1). Beyond speed breeding or indoor pollinations, embryo rescue can be employed to recover whole plants from field-pollinated seeds that have been impacted by disease or environmental damage, or would otherwise not be able to mature on the tree. For example, we have employed embryo rescue to recover important transgenic progeny from a wild-type hybrid mother tree that flowered outside of the typical pollination season. We performed controlled pollinations at the end of August, and embryo rescue in late September. Developing ovules would have typically been killed by the onset of cold temperatures, but embryo rescue allowed successful production of healthy plants from this late-season cross.

Third, combined methods of speed breeding and genetic analysis were aimed at identifying lab-grown homozygous transgenic American chestnut for pollen production to be crossed with wild-type mother trees in forests and germplasm conservation orchards located across the Eastern United States (Westbrook et al. 2020). We successfully recovered homozygous D54 transgenic lines from both fully matured and immature burs using speed breeding. At the time of writing, we have recovered a total of eleven homozygous D54 lines from 123 nuts or embryos (a 9% recovery rate) derived from crossing two hemizygous D54 parents bred in indoor, speed breeding settings (See Supplemental Table 3). Further, these methods led to the production of the first homozygous D54 transgenic line to produce viable pollen, which yielded 95% transgene inheritance in offspring from several mother trees. Pollen homozygous for *OxO* is a useful tool for research and breeding but is not seen as a critical component for restoration, since it may not enhance survival or reproduction of American chestnuts beyond the hemizygous state. Production and testing of additional ‘Darling 54’ homozygous lines, derived from tissue culture, full-term indoor breeding, and outdoor pollination, is underway to better understand crossing and inheritance rates with higher sample sizes from additional genetic backgrounds. While homozygous ‘Darling 54’ trees in this study demonstrated reproductive fitness in the form of rapid indoor growth and production of viable pollen, and anecdotal observations indicate typical phenotypes for young trees, metrics such as field growth and blight tolerance have not yet been compared to hemizygous trees. These methods can be easily adapted to produce and identify homozygous offspring of other transgenic events in the future.

In addition, this work highlights the value of integrating the methods of speed breeding, embryo rescue, and event-specific genotyping assays. We have validated two genotyping assays to determine copy number and to distinguish different transgenic lines. The qPCR assay provides a method to determine copy number on any transgenic events containing the *OxO* transgene, regardless of the promoter driving *OxO* or *a priori* knowledge of the genomic location for the transgenic event. This method can be helpful in early screening and characterization of transgenic lines, including improved lines such as those with wound-inducible promoters driving *OxO* expression (Carlson et al. 2022). On the other hand, performing the multiplex D58 and D54 assay on breeding material routinely can provide assurance to the speed breeding process, while also providing copy-number status.

The speed breeding and embryo rescue methods reported in this study are not restricted to transgenic chestnuts, although this method is especially important for controlled field pollinations under USDA-APHIS confined field trial permit conditions. Recent genomic advancements, such as the availability of high-quality genome assemblies of multiple *Castanea* species, facilitate research on transgenic American and traditional hybrid breeding work (Westbrook et al. 2025). Chestnut speed breeding as a methodology is relevant to all breeding-associated gains, such as trait development, genomic selection, backcross breeding, and genetic diversification efforts. Relatedly, Flowering Locus T (*FT*) transgenes have been engineered into trees including plum (Srinivasan et al. 2012) and poplar (Klocko et al. 2023) to directly address timing of flowering, which offers a potentially complementary approach for breeding and other applications. In addition, our embryo rescue methods can be utilized for conservation of unique genotypes, especially from threatened trees. It may also be useful to accelerate breeding of other *Castanea* species for agricultural purposes such as F1 *C. sativa* x *dentata* and *C. pumila* x *dentata* hybrids (Klak et al. 2021). This work highlights the continual improvements that can be made to rapid breeding of American chestnut and to rescuing immature embryos, both of which are critical for American chestnut research as we work towards the restoration of this iconic species back to its native range.

## Contributions and Acknowledgements

We gratefully acknowledge the foundational work of the late Drs. William Powell and Charles Maynard, without whom this research would not have been possible.

TK, HP, AN and EHT wrote the manuscript, and conceptualized the experimental design for speed breeding and molecular assays. TK, VGM, and HP performed speed breeding experiments and performed crosses to produce immature and mature seed. HP developed and tested embryo rescue protocols. TK, VGM, HP, and DM performed crosses to produce immature seeds, contributed to embryo rescue experiments, and provided DNA samples. DM performed and analyzed quantitative PCR. AO and EHT performed T1 mapping and junction analysis for ‘Darling 54.’ All authors reviewed the manuscript and provided edits.

Work by EHT was funded by USDA National Institute of Food and Agriculture, Hatch Project Number ME0-22405 through the Maine Agricultural & Forest Experiment Station, and NSF IOS award no. 2427904. Work by AN, HP, DM and AO was funded by the Templeton World Charity Foundation, the New York Chapter of The American Chestnut Foundation (now called American Chestnut Restoration, Inc.), and numerous individual donors. Work by TK and VGM was funded by the Quimby Family Foundation, the PW Sprague Memorial Foundation, The American Chestnut Foundation and its New York Chapter, and individual donors.

## Supplemental Data

Supplemental Figure 1: ‘Darling 54’ transgene insertion site on Chr04 of Ellis

Supplemental Figure 2: Images of mature and developing burs on sped bred American chestnut and histochemical assay from progeny from homozygous ‘Darling 54’ pollen

Supplemental Table 1: Oligo sequences and information on PCR products used in this study Supplemental Table 2: List of embryo rescue and fully mature chestnut seed used in this study Supplemental Table 3: Segregation ratios of progeny from two hemizygous ‘Darling 54’ parents from embryo cultures and mature seeds

## Notes

### Competing Interest Statement

The authors have declared no competing interest.

### Summary of Updates

Minor revisions following reviewer comments; additional data in Supplemental Table 3.

## References

Andrade GM, Nairn CJ, L. HT, Merkle SA. 2009. Sexually mature transgenic American chestnut trees via embryogenic suspension-based transformation. Plant Cell Rep. 28(9):1385–1397.

Bankevich A, Nurk S, Antipov D, Gurevich AA, Dvorkin M, Kulikov AS, Lesin VM, Nikolenko SI, Pham S, Prjibelski AD, et al. 2012. SPAdes: a new genome assembly algorithm and its applications to single-cell sequencing. J Comput Biol. 19(5):455–477.

Baier K, Maynard C, Powell W. 2012. Chestnut and light. The Journal of The American Chestnut Foundation. 26(3). https://tacf.org/wp-content/uploads/2016/09/Volume-XXVI-No.-3-May-June-2012.pdf

Bhatta M, Sandro P, Smith MR, Delaney O, Voss-Fels KP, Gutierrez L, Hickey LT. 2021. Need for speed: manipulating plant growth to accelerate breeding cycles. Curr Opin Plant Biol. 60:101986.

Burnham CR. 1988. The Restoration of the American Chestnut: Mendelian genetics may solve a problem that has resisted other approaches. Am Sci. 76(5):478–487.

Carlson E, Stewart K, Baier K, McGuigan L, Culpepper T 2nd, Powell W. 2022. Pathogen-induced expression of a blight tolerance transgene in American chestnut. Mol Plant Pathol. 23(3):370–382.

Castro CAO, dos Santos GA, Takahashi EK, Pires Nunes AC, Souza GA, & de Resende, MDV. 2021. Accelerating Eucalyptus breeding strategies through top grafting applied to young seedlings. Ind Crops Prod. 171:113906.

Cook R, Forest HS. 1979. The American chestnut II: Chestnuts in the Gennessee Valley region, 1978. The Rochester Committee for Scientific Inform Bul.

Flachowsky H, Le Roux P-M, Peil A, Patocchi A, Richter K, Hanke M-V. 2011. Application of a high-speed breeding technology to apple (Malus × domestica) based on transgenic early flowering plants and marker-assisted selection. New Phytol. 192(2):364–377.

Ghosh S, Watson A, Gonzalez-Navarro OE, Ramirez-Gonzalez RH, Yanes L, Mendoza-Suárez M, Simmonds J, Wells R, Rayner T, Green P, et al. 2018. Speed breeding in growth chambers and glasshouses for crop breeding and model plant research. Nat Protoc. 13(12):2944–2963.

Growing Chestnuts. [accessed 2025 Feb 20]. https://tacf.org/growing-chestnuts/.

Guilley H, Dudley RK, Jonard G, Balàzs E, Richards KE. 1982. Transcription of Cauliflower mosaic virus DNA: detection of promoter sequences, and characterization of transcripts. Cell. 30(3):763–773.

Hepting GH. 1974. Death of the American Chestnut. Journal of Forest History. 18(3):60–67.

Jiao Y, Li Z, Xu K, Guo Y, Zhang C, Li T, Jiang Y, Liu G, Xu Y. 2018. Study on improving plantlet development and embryo germination rates in in vitro embryo rescue of seedless grapevine. N Z J Crop Hortic Sci. 46(1):39–53.

Kaur R, Sharma N, Kumar K, Sharma DR, Sharma SD. 2006. In vitro germination of walnut (Juglans regia L.) embryos. Sci Hortic. 109(4):385–388.

Klak T, Spiers E, Powell WA. 2021. Transgenic pollen in less than a year. Chestnut: The Journal of The American Chestnut Foundation 35(3):27–29.

Klocko AL, Elorriaga E, Ma C, Strauss SH. Variation in floral form of CRISPR knock-outs of the poplar homologs of LEAFY and AGAMOUS after FT heat-induced early flowering. Hort Res. 10:uhad132.

Kolpak SE, Smith J, Albrecht MJ, DeBell J, Lipow S, Cherry ML, Howe GT. 2015. High-density miniaturized seed orchards of Douglas-fir. New For. 46:121–140.

Kyte L, Kleyn J, Scoggins H, Bridgen M. 2013. Plants from Test Tubes: An Introduction to Micropropagation. Timber Press.

Liang H, Maynard CA, Allen RD, Powell WA. 2001. Increased Septoria musiva resistance in transgenic hybrid poplar leaves expressing a wheat oxalate oxidase gene. Plant Mol Biol. 45(6):619–629.

Maynard CA, McGuigan LD, Oakes AD, Zhang B, Newhouse AE, Northern LC, Chartrand AM, Will LR, Baier KM, Powell WA. 2015. Chestnut, American (Castanea dentata (Marsh.) Borkh.). Methods Mol Biol. 1224:143–161.

McGuigan L, Fernandes P, Oakes A, Stewart K, Powell W. 2020. Transformation of American chestnut (Castanea dentata (Marsh.) Borkh) using RITA® temporary immersion bioreactors and We Vitro containers. Forests. 11(11):1196.

McKay J. 1942. Self-sterility in the Chinese chestnut (Castanea mollissima). Proc Am Soc Hort Sci. 41:156–160.

Moore GA, Rogers KL, Kamps TL, Pajon M, Stover J, Febres VJ, Stover E. 2016. Rapid cycling plant breeding in citrus. Citrograph 7(3):80–85.

Newhouse AE, Polin-McGuigan LD, Baier KA, Valletta KER, Rottmann WH, Tschaplinski TJ, Maynard CA, Powell WA. 2014. Transgenic American chestnuts show enhanced blight resistance and transmit the trait to T1 progeny. Plant Sci. 228:88–97.

Newhouse AE, Powell WA. 2021. Intentional introgression of a blight tolerance transgene to rescue the remnant population of American chestnut. Conserv Sci Pract. 3(4):e348.

Newhouse AE. 2025. Petition for determination of nonregulated status for blight-tolerant Darling 54 American chestnut. [accessed 2025 Oct 14]. https://www.aphis.usda.gov/sites/default/files/19-30901p-rev.pdf

van Nocker S, Gardiner SE. 2014. Breeding better cultivars, faster: applications of new technologies for the rapid deployment of superior horticultural tree crops. Hortic Res. 1:14022.

Oakes AD, Pilkey HC, Powell WA. 2020. Improving ex vitro rooting and acclimatization techniques for micropropagated American Chestnut. J Environ Hortic. 38(4):149–157.

Paillet FL, Rutter PA. 1989. Replacement of native oak and hickory tree species by the introduced American chestnut (Castanea dentata) in southwestern Wisconsin. Can J Bot. 67(12):3457–3469.

Pilkey H. 2021. Developing and Optimizing Outcrossing Conservation Methods to Aid Diversification and Production of Blight-Tolerant American Chestnut Trees (Castanea dentata [Marsh.] Borkh.). SUNY College of Environmental Science and Forestry. [accessed 2025 Apr 3]. https://experts.esf.edu/esploro/outputs/99895578804826.

Powell WA, Newhouse AE, Coffey V. 2019. Developing Blight-Tolerant American Chestnut Trees. Cold Spring Harb Perspect Biol. 11(7). doi:10.1101/cshperspect.a034587. http://dx.doi.org/10.1101/cshperspect.a034587.

Srinivasan C, Dardick C, Callahan A, Scorza R. 2012. Plum (Prunus domestica) trees transformed with poplar FT1 result in altered architecture, dormancy requirement, and continuous flowering. PLoS One 7(7):e40715.

Steiner KC, Westbrook JW, Hebard FV, Georgi LL, Powell WA, Fitzsimmons SF. 2017. Rescue of American chestnut with extraspecific genes following its destruction by a naturalized pathogen. New Forests. 48(2):317–336.

Stilwell KL, Wilbur HM, Werth CR, Taylor DR. 2003. Heterozygote advantage in the American chestnut, Castanea dentata. Am J Bot. 90(2):207–213.

Watson A, Ghosh S, Williams MJ, Cuddy WS, Simmonds J, Rey M-D, Asyraf Md Hatta M, Hinchliffe A, Steed A, Reynolds D, et al. 2018. Speed breeding is a powerful tool to accelerate crop research and breeding. Nat Plants. 4(1):23–29.

Westbrook JW, Holliday JA, Newhouse AE, Powell WA. 2020. A plan to diversify a transgenic blight‐tolerant American chestnut population using citizen science. Plants People Planet. 2(1):84–95.

Westbrook JW, Malukiewicz J, Sreedasyam A, Jenkins JW, Zhang Q, Lakoba V, Fitzsimmons SF, Van Clief J, Collins K, Hoy S, et al. 2025. Improving American chestnut resistance to two invasive pathogens through genome-enabled breeding. bioRxiv.:2025.01.30.635736. doi:10.1101/2025.01.30.635736. [accessed 2025 Feb 6]. https://www.biorxiv.org/content/10.1101/2025.01.30.635736v1.abstract.

Xing Z, Powell WA, Maynard CA. 1999. Development and germination of American chestnut somatic embryos. Plant Cell Tissue Organ Cult. 57(1):47–55.

Yang Y, Zhang D, Li Z, Jin X, Dong J. 2015. Immature Embryo Germination and Its Micropropagation of Ilex crenata Thunb. HortScience. 50(5):733–737.

Zhang B, Oakes AD, Newhouse AE, Baier KM, Maynard CA, Powell WA. 2013. A threshold level of oxalate oxidase transgene expression reduces Cryphonectria parasitica-induced necrosis in a transgenic American chestnut (Castanea dentata) leaf bioassay. Transgenic Res. 22(5):973–982.

