## Supplemental Figures and Tables for "Speed breeding transgenic American chestnut trees toward restoration"

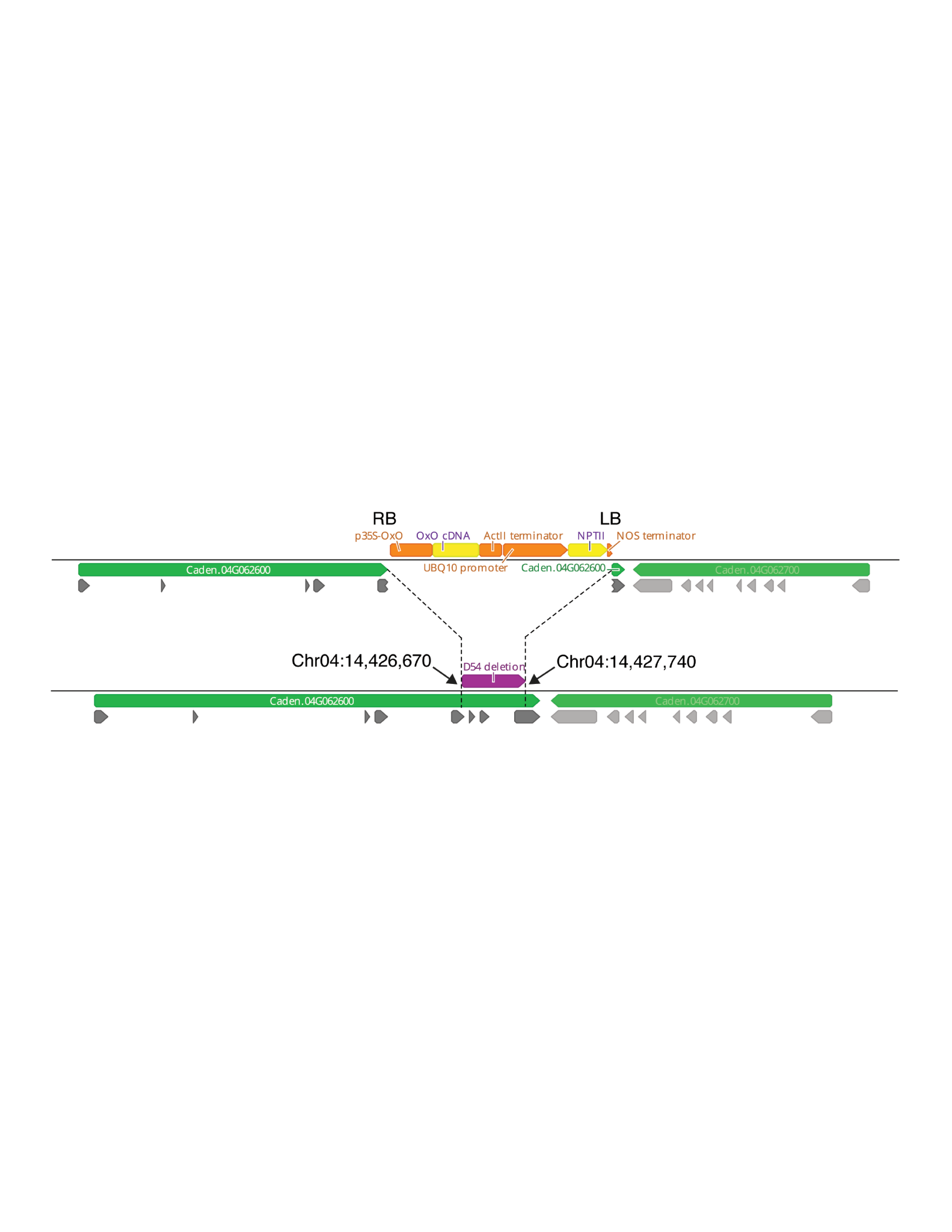

Supplemental Figure 1. **‘Darling 54’ transgene insertion site on Chr04 of Ellis.** Reconstruction of the ‘Darling 54’ insertion site on Chr04 based on the Ellis1 v1.1 genome assembly. The *35S::OxO* T-DNA for ‘Darling 54’ is inserted between Chr04:14,426,670 (RB junction) and Chr04:14,427,740 (LB junction), resulting in a 1,069 bp deletion. This event partially deleted exon 5 and exon 8 of Caden.04G062600 and the intervening sequences. Dark gray shaded boxes are exons in the direction of its open reading frame for Caden.04G062600 gene model and neighboring gene, Caden.04G062700 is 186 bp away and encoded in the opposite direction (light gray shaded boxes). Caden.04G062600 appears to be the orthologous *cdSAL1* gene in American chestnut.

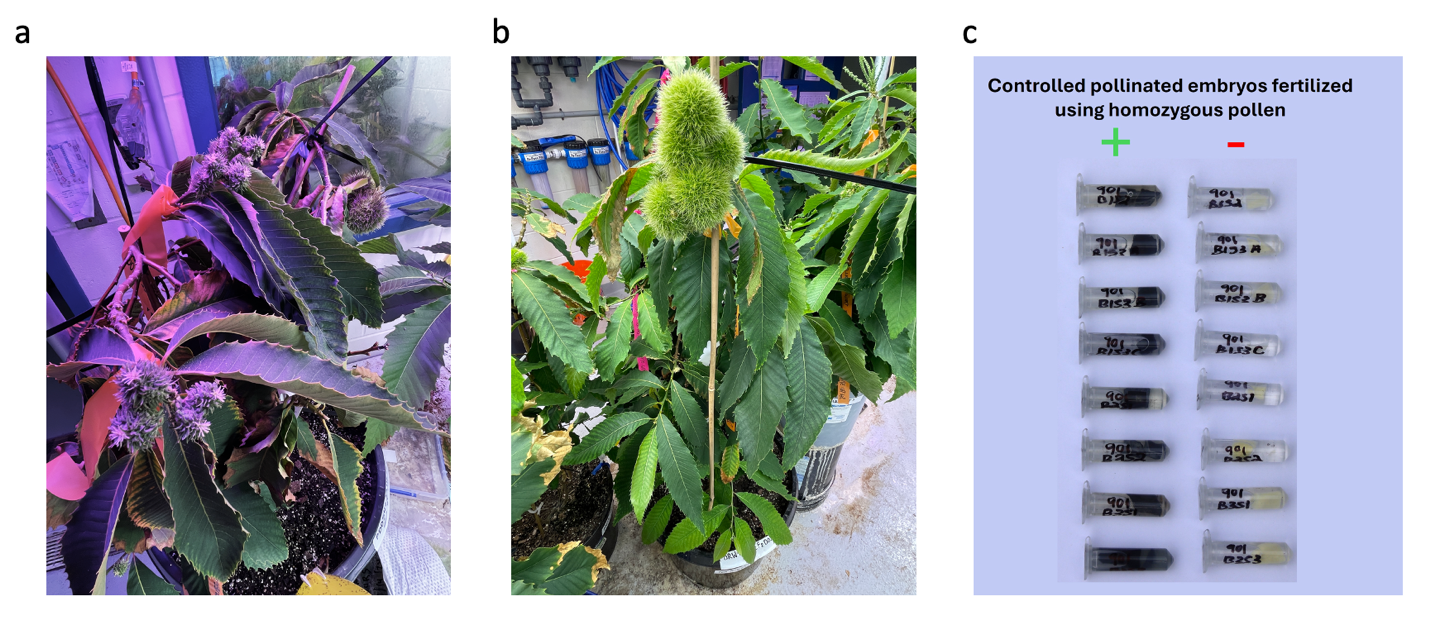

Supplemental Figure 2. **Images of actively developing and mature burs on sped bred transgenic American chestnut and histochemical assay from progeny from sped bred homozygous ‘Darling 54.’** (a) Actively developing burs and (b) mature burs on sped bred transgenic American chestnut. (c) Histochemical assay of progeny derived from controlled pollinations using homozygous ‘Darling 54’ (D54) pollen from TGA002 (+) compared to negative wild-type control pollination (-) showing presence of D54 transgene in all embryos from TGA002 crosses.

Supplemental Table 1. **Oligo sequences and information on PCR products used in this study.**

| **Oligo ID** | **Oligo Sequence** |  |
| --- | --- | --- |
| qPCR_OxOF | CAGCGGCAAACTTGGACTTGAGAA |  |
| qPCR_OxOR | TGCACTTCCAGTTCAACGTCGGTA |  |
| Actin F | CCTTGCTGGTCGTGATCTC |  |
| Actin R | GTCTCAAGTTCCTGGCTCATAGTC |  |
| AC-UBQ F | GAGTCCACATCCACCTCGT |  |
| AC-UBQ R | CCTCCAAGGTGATGGTTTTG |  |
| SX58_chr7.5 | CGTTATATGTTACACCACCCAATAAC |  |
| 35S:OxO_RB3 | CTTGCTAGCTTCTGCAGGTACCT |  |
| SX58_chr7.4 | CTACTTGCGTGCACGGTTTC |  |
| 35S:OxO_LB1 | GCCTTCTATCGCCTTCTTG |  |
| SX58_3R | GCGACCAAATTAGTCACAACTTAGAAGTGG |  |
| D54_Chr4.1 | GCTATGGTGACATCTCAGGTGTTTGGCA |  |
| D54_Chr4.7 | CCTGCCTTTCTTGGCTTCAAGT |  |
| D54_Chr4.5 | GGATTCAAAGAGTGATGCTTCCTCTGG |  |
| **PCR Name** | **Oligo Combinations** | **Expected Sizes (bp)** |
| OxO qPCR | qPCR_OxOF + qPCR_OxOR | 192 |
| D58 RB | 35S:OxO_RB3 + SX58_chr7.5 | 411 |
| D58 LB | 35S:OxO_LB1 + SX58_chr7.4 | 391 |
| D58 Multiplex | 35S:OxO_LB1 + SX58_chr7.4 + SX58_3R | 566 + 391 |
| D54 RB | 35S:OxO_RB3 + D54_Chr4.1 | 479 |
| D54 LB | 35S:OxO_LB1 + D54_Chr4.7 | 854 |
| D54 Multiplex | 35S:OxO_RB3 + D54_Chr4.1 + D54_Chr4.5 | 599 + 479 |

Supplemental Table 2. **List of embryo rescue and fully mature chestnut seed used in this study.** Histochemical assay (“+” = OxO activity, “-” = no activity). Multiplex PCR distinguishes lines that are wild-type (wt) having no ‘Darling 54’ transgene, hemizygous or homozygous for *OxO*. OxO qPCR assay is based on relative copy number of *OxO* to housekeeping genes (0 copy considered wt, 1 copy considered het and 2 copies considered homozygous). *UERb001 was considered wild-type from the absence of OxO activity by histochemical staining.

| **Line ID** | **Source** | **Histochemical** | **Multiplex PCR** | **OxO qPCR** | **Generation** |
| --- | --- | --- | --- | --- | --- |
| TGA001 | Embryo rescue | - | not tested | 0 copy | T3 |
| TGA002 | Embryo rescue | + | homozygous | 2 copies | T3 |
| TGE001 | Embryo rescue | + | not tested | 1 copy | T5 |
| TGF001 | Embryo rescue | + | not tested | 1 copy | T5 |
| TGF011 | Embryo rescue | + | not tested | 1 copy | T5 |
| TGF012 | Embryo rescue | + | not tested | 1 copy | T5 |
| TGF013 | Embryo rescue | + | not tested | 1 copy | T5 |
| TGG001 | Embryo rescue | + | not tested | 1 copy | T5 |
| TGG002 | Embryo rescue | + | not tested | 2 copies | T4 |
| TGG003 | Embryo rescue | - | not tested | 0 copy | T4 |
| TGG004 | Embryo rescue | + | homozygous | 2 copies | T4 |
| TGG008 | Embryo rescue | + | hemizygous | 1 copy | T4 |
| TGG009 | Embryo rescue | + | hemizygous | 1 copy | T4 |
| TGG010 | Embryo rescue | - | wt | 0 copy | T4 |
| TGH001 | Embryo rescue | + | not tested | 1 copy | T5 |
| TGH002 | Embryo rescue | + | not tested | 1 copy | T5 |
| TGH004 | Embryo rescue | - | not tested | 0 copy | T5 |
| TGH005 | Embryo rescue | + | not tested | 1 copy | T5 |
| TGJ001 | Embryo rescue | + | not tested | 1 copy | T5 |
| TGJ002 | Embryo rescue | + | not tested | 1 copy | T5 |
| TGJ003 | Embryo rescue | + | not tested | 1 copy | T5 |
| TGJ004 | Embryo rescue | - | not tested | 0 copy | T5 |
| TGJ005 | Embryo rescue | + | hemizygous | 1 copy | T5 |
| TGJ006 | Embryo rescue | + | not tested | 1 copy | T5 |
| GF005 | Mature seedling | + | hemizygous | not tested | T4 |
| GF006 | Mature seedling | - | wt | not tested | T4 |
| GF014 | Mature seedling | not tested | hemizygous | not tested | T4 |
| LA001 | Mature seedling | + | hemizygous | not tested | T5 |
| LA002 | Mature seedling | - | wt | not tested | T5 |
| LA003 | Mature seedling | + | hemizygous | not tested | T4 |
| UEFe004 | Mature seedling | + | hemizygous | not tested | T5 |
| UEFe005 | Mature seedling | + | hemizygous | not tested | T5 |
| UEFe006 | Mature seedling | + | hemizygous | not tested | T5 |
| UEFe007 | Mature seedling | - | wt | not tested | T5 |
| UEFe008 | Mature seedling | - | wt | not tested | T5 |
| UEFe009 | Mature seedling | - | wt | not tested | T5 |
| UEFe012 | Mature seedling | + | hemizygous | not tested | T5 |
| *UERb001 | Mature seedling | - | pcr failure | not tested | T4 |
| UERb003 | Mature seedling | - | wt | not tested | T4 |
| UERb004 | Mature seedling | + | hemizygous | not tested | T4 |
| UERb005 | Mature seedling | + | hemizygous | not tested | T5 |
| UERb006 | Mature seedling | - | wt | not tested | T5 |
| UERb013 | Mature seedling | + | hemizygous | not tested | T4 |
| UERb016 | Mature seedling | - | wt | not tested | T4 |
| UERb017 | Mature seedling | not tested | hemizygous | not tested | T4 |
| ULB002 | Mature seedling | - | wt | not tested | T4 |
| ULL002 | Mature seedling | - | wt | not tested | T5 |
| ULL005 | Mature seedling | not tested | hemizygous | not tested | T5 |
| ULL009 | Mature seedling | not tested | homozygous | not tested | T5 |
| ULL010 | Mature seedling | not tested | wt | not tested | T5 |
| ZX001 | Mature seedling | - | wt | not tested | T4 |
| ZX002 | Mature seedling | + | hemizygous | not tested | T4 |
| ZX003 | Mature seedling | + | hemizygous | not tested | T4 |
| ZX004 | Mature seedling | + | hemizygous | not tested | T4 |
| ZX005 | Mature seedling | + | hemizygous | not tested | T4 |
| ZX007 | Mature seedling | + | hemizygous | not tested | T4 |
| ZX008 | Mature seedling | - | wt | not tested | T4 |
| ZX009 | Mature seedling | + | hemizygous | not tested | T4 |
| ZX010 | Mature seedling | - | wt | not tested | T4 |
| ZX011 | Mature seedling | + | hemizygous | not tested | T4 |
| ZX012 | Mature seedling | - | wt | not tested | T4 |

Supplemental Table 3. **Segregation ratios of progeny from two hemizygous ‘Darling 54’ (D54) parents from embryo culture of immature seeds and from fully mature seeds.** Number of offspring observed to be homozygous D54 (HM), hemizygous D54 (HEM) or wild-type (WT) are listed along with p-value of each 𝜒^2^ test based on each contingency table. In total, eleven homozygous individuals were identified from 123 embryos/seeds.

| Embryo Culture | HM | HEM | WT |  | Mature Seed | HM | HEM | WT |
| --- | --- | --- | --- | --- | --- | --- | --- | --- |
| Observed | 3 | 16 | 5 |  | Observed | 8 | 64 | 27 |
| Expected | 6 | 1212 | 6 |  | Expected | 24.75 | 49.5 | 24.75 |
| 𝜒^2^ *p*-value | 0.22313016 |  |  |  | 𝜒^2^ *p*-value | 0.000373 |  |  |
